# A plasmid-encoded H-NS protein selectively binds its own plasmid

**DOI:** 10.64898/2026.06.14.732234

**Authors:** Anne M. Stringer, Diana Rodríguez-Valverde, Fernando Ruiz-Perez, Araceli E. Santiago, Joseph T. Wade

## Abstract

H-NS is an abundant nucleoid-associated protein found in *Enterobacterales* species. Some conjugative plasmids encode H-NS homologues, which are thought to facilitate plasmid maintenance and reduce the fitness costs associated with plasmid carriage. Here, we characterize HppX_CROD2_, an H-NS homologue encoded by the IncX4 plasmid pCROD2 of *Citrobacter rodentium*. Our data indicate that HppX_CROD2_ has a strong preference for binding pCROD2 over the chromosome or other plasmids. By contrast, chromosomally encoded H-NS displays no preference for plasmid sequence. When expressed from a heterologous plasmid in *Escherichia coli*, HppX_CROD2_ showed similar DNA-sequence preference to chromosomally encoded H-NS. Moreover, HppX_CROD2_ binding to a sequence from pCROD2 was much lower when that sequence was cloned in a laboratory plasmid. Thus, HppX_CROD2_ preferentially binds DNA in the context of the plasmid where it is encoded, a phenomenon we term “cognate plasmid specificity”. We propose that cognate plasmid specificity occurs through recognition of plasmid-specific DNA topology generated by plasmid-encoded topoisomerases. Cognate plasmid specificity may insulate regulation of plasmid genes from the effects of host DNA, while minimizing disruption of host chromosome regulation due to plasmid carriage.

**IMPORTANCE:** Many bacteria carry conjugative plasmids, mobile DNA molecules that spread traits such as antibiotic resistance. Some conjugative plasmids encode proteins related to the bacterial DNA-binding protein H-NS. We show that an H-NS-like protein from the IncX4 plasmid pCROD2 binds almost exclusively to the plasmid from which it originates, while largely ignoring the host chromosome. Our findings reveal a previously unrecognized mechanism that allows plasmids to regulate their own genes with high specificity while minimizing interference with host gene expression.

## MAIN TEXT

“<u>N</u>ucleoid-<u>A</u>ssociated <u>P</u>roteins” (NAPs) are a loosely defined group of abundant DNA-binding proteins in bacteria that bind DNA with low sequence specificity (1). The best-studied nucleoid-associated protein is the <u>H</u>istone-like <u>N</u>ucleoid-<u>S</u>tructuring protein (H-NS), which is found primarily in *Enterobacterales* species (2). H-NS binds A/T-rich DNA, can nucleate along stretches of A/T-rich sequence, and can form bridges between distal DNA regions (2). In *Escherichia coli* and *Salmonella*, H-NS binds ∼10% of the chromosome, predominantly in horizontally acquired sequence (3–7). Binding of H-NS silences transcription of A/T-rich genes (3, 6), and silences transcription from cryptic promoters found within A/T-rich genes (8). Loss of H-NS derepresses large numbers of promoters, leading to the titration of RNA polymerase away from housekeeping genes (9).

Conjugative plasmids often encode NAP homologues (1). Plasmid-encoded H-NS homologues, termed “Hpp” proteins, are encoded on diverse conjugative plasmids (10). Like the canonical, chromosomally encoded H-NS (referred to here as H-NS_chr_), Hpp proteins bind DNA with a strong preference for A/T-rich DNA, leading to considerable overlap in the chromosomal distribution of H-NS_chr_ and Hpp binding (11–14). Hpp proteins have been shown to repress expression of DNA transfer genes (12, 13, 15–17), promote plasmid stability (16, 18), and modulate fitness of their bacterial hosts (15, 16, 18, 19). Sfh, an Hpp protein encoded on an IncHI plasmid, promotes bacterial fitness by supplementing H-NS_chr_, thereby preventing titration of H-NS_chr_ away from chromosomal sites by the A/T-rich plasmid. Thus, it has been proposed that Hpp proteins provide a “stealth” function that insulates their cognate plasmid from the bacterial host (20).

The pCROD2 plasmid from *Citrobacter rodentium* is from the IncX4 family and encodes an Hpp that we term HppX_CROD2_ according to a previously described classification system (Figure 1A) (10). We compared the sequence of HppX_CROD2_ to the *C. rodentium* H-NS_chr_ and several characterized Hpp proteins encoded on other plasmids (Figure S1). HppX_CROD2_ has between 19% and 45% sequence identity with the other plasmid-encoded Hpp proteins, and 37% sequence identity with H-NS_chr_. Thus, the sequence of HppX_CROD2_ differs substantially from that of other characterized Hpp proteins.

**Figure 1.**
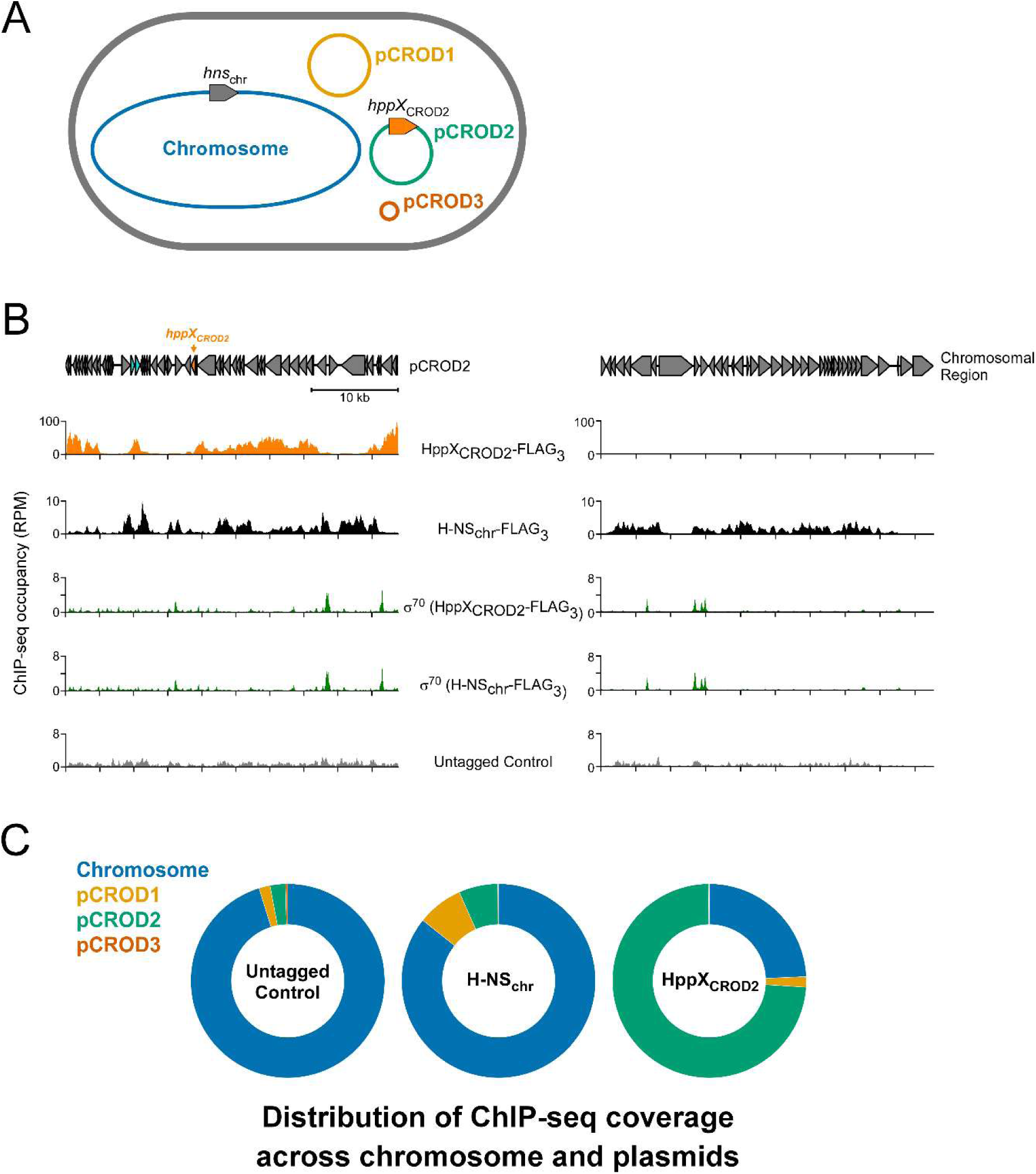
HppX_CROD2_ binds selectively to the pCROD2 plasmid in *C. rodentium*. **(A)** Schematic of the *C. rodentium* DBS100 strain showing the genomic locations of *hns_chr_* and *hppX_CROD2_*. **(B)** ChIP-seq coverage for HppX_CROD2_, H-NS_chr_, and σ^70^ across pCROD2 (left panel) and a similarly sized region of the chromosome (genome positions 4,350,200-4,388,300; right panel). Note that σ^70^ ChIP-seq data are shown for each of the HppX_CROD2_-FLAG_3_ and H-NS_chr_-FLAG_3_ strains. One of two replicate datasets is shown for each experiment. **(C)** Donut-charts showing the proportion of ChIP-seq coverage mapping to the *C. rodentium* chromosome and plasmids for an untagged control, H-NS_chr_-FLAG_3_, and HppX_CROD2_-FLAG_3_.

To determine which DNA regions are bound by HppX_CROD2_, we used ChIP-seq to map the genome-wide association of a C-terminally FLAG_3_-tagged derivative of HppX_CROD2_ across the *C. rodentium* genome, which includes the chromosome, pCROD2, and two other plasmids. In parallel, we used ChIP-seq to map the genome-wide association a C-terminally FLAG_3_-tagged derivative of H-NS_chr_. We observed association of HppX_CROD2_ with regions across the pCROD2 plasmid (Figure 1B), consistent with the plasmid’s relatively high A/T-content (58%). However, we observed little association of HppX_CROD2_ with the chromosome or with the other plasmids. By contrast, H-NS_chr_ was associated with many regions across the chromosome (Figure 1B), predominantly in areas of high A/T-content, as expected (Figure S2). While we did observe H-NS_chr_ association with pCROD2, H-NS_chr_ binding was largely limited to regions not bound by HppX_CROD2_ (Figure 1B). When comparing the binding of HppX_CROD2_ and H-NS_chr_, we observed a ∼40-fold higher specificity of HppX_CROD2_ for pCROD2 than we observed for H-NS_chr_ (Figure 1C). We conclude that HppX_CROD2_ is highly selective for binding pCROD2. We considered the unlikely possibility that epitope-tagging altered the function of HppX_CROD2_ in a way that increased pCROD2 copy number, which could explain higher HppX_CROD2_ ChIP-seq coverage on pCROD2. To test this idea, we used ChIP-seq to map the genome-wide association of σ^70^ in each of the HppX_CROD2_-FLAG_3_ and H-NS_chr_-FLAG_3_ strains. We observed very similar σ^70^ association with both the chromosome and pCROD2 in the two strains (Figure 1B; R^2^ values of 0.84-0.91 when comparing datasets from the two strain backgrounds). We conclude that tagging HppX_CROD2_ does not affect pCROD2 copy number.

We reasoned that the strong preference of HppX_CROD2_ for pCROD2 could be due to the presence of optimal HppX_CROD2_ binding sites on pCROD2 that are missing from the chromosome, although we thought this unlikely given the relative sizes of pCROD2 and the chromosome. Nonetheless, to test this hypothesis, we cloned a ∼1.2 kb bp region from pCROD2 that has high HppX_CROD2_ ChIP-seq signal, into a laboratory cloning plasmid, pAMD407. We then introduced pAMD407 into *E. coli* K-12 together with a separate laboratory cloning plasmid (pAMD406) encoding HppX_CROD2_-FLAG_3_ (Figure 2A). We used ChIP-seq to map the association of HppX_CROD2_ with the *E. coli* chromosome and the two plasmids. The profile of HppX_CROD2_ binding across the ∼1.2 kb pCROD2 sequence closely mirrored that observed on pCROD2 itself in *C. rodentium*. However, the level of HppX_CROD2_ binding to this region was comparable to regions on the chromosome (Figure 2B-C). Moreover, HppX_CROD2_ binding across the *E. coli* chromosome was similar to that observed previously for H-NS_chr_ (13) (Figure S3A; H-NS_chr_ from *E. coli* is 96% identical to that from *C. rodentium*), albeit with a slightly reduced requirement for A/T-richness (Figure S3B). HppX_CROD2_ binding across the *E. coli* chromosome was also higher than the binding we observed across the *C. rodentium* chromosome, perhaps because of titration of HppX_CROD2_ by pCROD2. We conclude that the strong preference of HppX_CROD2_ for pCROD2 is not simply due to its DNA sequence specificity. Additionally, because we observed little association of HppX_CROD2_ with pAMD406, the plasmid where it was encoded in this experiment (Figure S4), we conclude that HppX_CROD2_ has a strong preference for pCROD2 specifically, rather than any plasmid where it is encoded.

**Figure 2.**
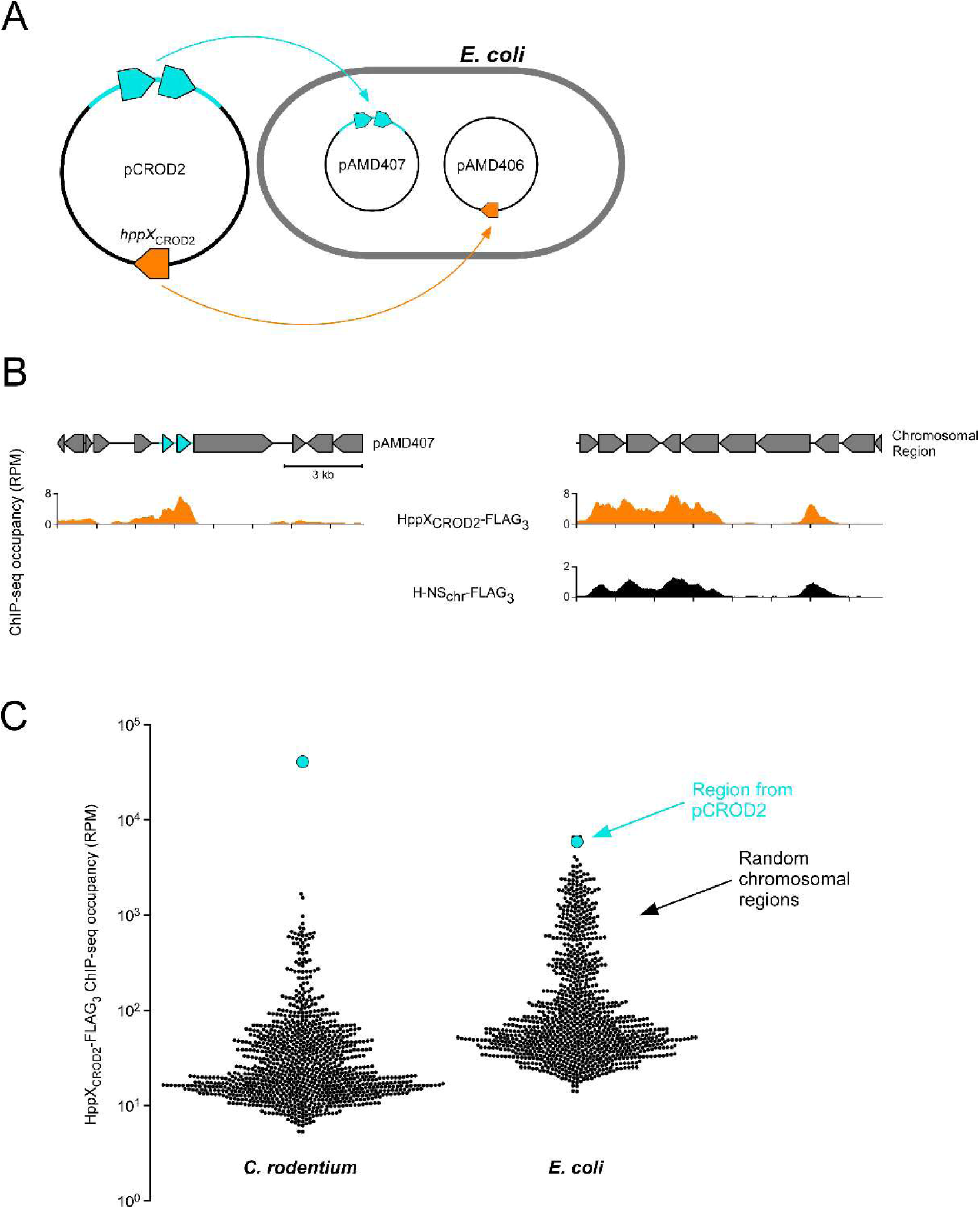
HppX_CROD2_ binding to pCROD2 sequence is greatly reduced outside the context of pCROD2. **(A)** Schematic showing the cloning plasmids introduced into *E. coli*. **(B)** ChIP-seq coverage for HppX_CROD2_ (left and right panels) and H-NS_chr_ (the *E. coli* protein; right panel only) across a plasmid containing a region of pCROD2 (shown in cyan), and a similarly sized region of the *E. coli* chromosome. Note that ChIP-seq data for H-NS_chr_ are from a strain lacking HppX_CROD2_. One of two replicate datasets is shown for each experiment. **(C)** Swarm plot showing HppX_CROD2_-FLAG_3_ ChIP-seq coverage across 1,000 randomly selected chromosomal regions and the pCROD2 region on a heterologous plasmid in *E. coli*.

During the course of this work, Wang *et al*. reported a ChIP-seq analysis of the Hpp protein Sfx, encoded on the R6K plasmid (IncX2 family) in an *E. coli* host. Similar to our observation for HppX_CROD2_, Wang *et al*. observed a strong preference for Sfx to bind the R6K plasmid over the chromosome, although they suggest that Sfx selectively binds only a portion of R6K. Moreover, Gao *et al*, recently reported a ChIP-seq analysis of an Hpp protein, HppX3, that is similar to Sfx (77% amino acid identity). HppX3 is encoded on the *bla*_NDM-5_-IncX3 plasmid. Our analysis of HppX3 ChIP-seq data from *E. coli* revealed a ∼6-fold enrichment of HppX3 binding to *bla*_NDM-5_-IncX3 relative to the chromosome (Figure S5). This enrichment is likely due to specific recognition of the plasmid rather than DNA sequence specificity because the A/T-content of the plasmid is relatively low (53%). Thus, three different Hpp proteins from three different families of IncX plasmids appear to have a preference for binding the plasmid on which they are natively encoded. We refer to this phenomenon as “cognate plasmid specificity”.

What is the basis of cognate plasmid specificity for Sfx, HppX3, and HppX_CROD2_? Wang *et al*. showed that treating *E. coli* with novobiocin, a topoisomerase inhibitor, prevents Sfx-mediated repression of R6K conjugation genes. Intriguingly, SfX, HppX3, and HppX_CROD2_ are all encoded either immediately adjacent to, or one gene away from a topoisomerase gene (Figure S6). We speculate that the plasmid-encoded topoisomerase specifically targets the plasmid to generate a distinct DNA topology that drives HppX cognate plasmid specificity.

Cognate plasmid specificity may provide fitness advantages for both the plasmid and its bacterial host. The same plasmid can reside in distantly related bacterial hosts, potentially with different amounts of A/T-rich sequence on the chromosome and on other plasmids. Cognate plasmid specificity would insulate the Hpp protein from these external sequences, ensuring appropriate repression of conjugation genes in any bacterial host. Loss of cognate plasmid specificity may lead to uncontrolled expression of conjugation genes that would negatively impact fitness due to the high energetic cost of producing the DNA transfer machinery (21) and exposure of the bacterial host to infection by plasmid-dependent phage that bind conjugative pili (22). Cognate plasmid specificity would also limit the impact of plasmid carriage on the bacterial host by preventing H-NS_chr_ binding to the plasmid (20). Titration of H-NS_chr_ away from chromosomal targets would reduce bacterial fitness by reducing transcription of housekeeping genes (9). Consistent with this idea, chromosomally encoded H-NS appears to be excluded from plasmid DNA by HppX_CROD2_ and Sfx (Figure 1A and (13)).

## SUPPLEMENTARY MATERIALS AND METHODS

### Strains and plasmids

All plasmids and strains used in this study are listed in Table S1. All oligonucleotides, and chemically synthesized double-stranded DNA fragments used in this study are listed in Table S2.

*C. rodentium* strains DBS100 (wild-type), JPCR7 (H-NS_chr_-FLAG_3_), and JPCR9 (HppX_CROD2_-FLAG_3_) have been described previously (23, 24). *E. coli* strain MG1655 has been described previously (25).

Plasmid pAMD403 was constructed by PCR-amplification of DNA fragment GB30 using oligonucleotides JW12508 and JW12509. GB30 encodes C-terminally FLAG_3_-tagged HppX_CROD2_ under the control of a constitutive promoter (26). The resulting PCR product was cloned into plasmid pBAD33 (27) digested with *Cla*I and *Hin*dIII. All screened clones contained mutations predicted to weaken either the Shine-Dalgarno or promoter activity. Plasmid pAMD403 contains a –12 (with respect to the start codon of *hppX_CROD2_*) G to T substitution, a change that is expected to weaken the Shine-Dalgarno sequence. Plasmid pAMD406 was constructed by PCR-amplifying HppX_CROD2_ –FLAGX_3_ together with the promoter and 5’ UTR from pAMD403 using oligonucleotides JW12508 and JW13094. The resulting PCR product was cloned into plasmid pBAD30 (27) digested with *Cla*I and *Sac*I. Plasmid pAMD407 was constructed by PCR-amplifying DNA fragment GB49 with oligonucleotides JW13223 and JW13224. GB49 contains a 1,268 bp sequence from the pCROD2 plasmid. The resulting PCR product was cloned into pAMD-BA-*lacZ* (28) digested with *Sph*I and *Hin*dIII.

### ChIP-seq

For ChIP-seq experiments with *C. rodentium*, strains JPCR7, JPCR9, and DBS100 were grown overnight in LB medium and diluted 1:70 into Dulbecco’s Modified Eagle Medium. The JPCR7 and JPCR9 cultures were supplemented with kanamycin (50 μg/mL). Cultures were grown to an OD_600_ of 0.7-0.8. For ChIP-seq experiments with *E. coli* MG1655 harboring pAMD406 and pAMD407, overnight cultures were diluted 1:100 in LB medium supplemented with ampicillin (100 μg/mL) and chloramphenicol (30 μg/mL), and grown to an OD600 of ∼0.6. All cells were processed for ChIP-seq as described previously (28) using 2 μl M2 anti-FLAG antibody (Sigma-Aldrich) or 2 μl anti-*E. coli* σ^70^ (BioLegend). All ChIP-seq experiments were performed in biological duplicate. R^2^ values for replicate pairs were 0.66 (strain DBS100; *C. rodentium* untagged control), 0.95 (strain JPCR7; H-NS_chr_ in *C. rodentium*), 0.99 (strain JPCR9; HppX_CROD2_ in *C. rodentium*), 0.84 (σ^70^ for the H-NS_chr_-FLAG_3_ strain of *C. rodentium*), 0.93 (σ^70^ for the HppX_CROD2_-FLAG_3_ strain of *C. rodentium*), and 0.98 (HppX_CROD2_ in *E. coli*).

### Analysis of ChIP-seq data

*C. rodentium* ChIP-seq paired-end sequence reads were aligned using Rockhopper (version 2.03) to the *C. rodentium* ICC168 reference genome (GenBank assembly GCA_000027085.1) in which the chromosome and plasmids were concatenated into a single reference. For ChIP-seq of HppX_CROD2_ in *E. coli*, paired-end sequence reads were aligned using Rockhopper to a reference genome in which the *E. coli* MG1655 chromosome (GenBank ID U00096.3) and plasmids pAMD407 and pAMD406 were concatenated into a single reference. Output .wig files were converted to .gff format using a custom Python script (version 3.12.0). All additional analyses used the.gff format to determine sequence read coverage per genome position. H-NS_chr_ ChIP-seq data for *E. coli* have been published previously (13), as have ChIP-seq data for HppX3 together with corresponding data for input sample (12). All additional analyses, and creation of genome browser images, used custom Python code. When determining the distribution of sequence reads across different genome elements, the first and last 500 bp of each genomic element were excluded.

Swarm plots showing HppX_CROD2_-FLAG_3_ ChIP-seq coverage were generated by summing normalized ChIP-seq coverage (reads per million) for a 1,267 bp on pCROD2, either in the context of pCROD2 in *C. rodentium*, or cloned into pAMD407 in *E. coli*. Normalized ChIP-seq coverage (reads per million) was also summed for 1,000 randomly selected chromosomal regions of 1,267 bp.

For the scatter-plot showing H-NS_chr_ and HppX_CROD2_ ChIP-seq occupancy, normalized ChIP-seq coverage (reads per million) was summed for 10,000 randomly selected 100 bp regions of the *E. coli* chromosome.

For analysis of the relationship between ChIP-seq signal and A/T content, normalized ChIP-seq occupancy (reads per million) was summed across 1,000,000 randomly selected 101 bp regions. A/T content was determined for the same regions. Mean ChIP-seq occupancy was calculated for regions sorted into bins based on A/T content, with each bin spanning 1%.

### Protein sequence alignment

Protein sequences were aligned using Clustal Omega (29) and visualized using Jalview (30).

### Plasmid alignment

The R6K, *bla*_NDM-5_-IncX3, and pCROD2 plasmids were aligned using clinker (31).

## DATA AVAILABILITY

Raw and processed ChIP-seq data are available at EBI ArrayExpress using accession number E-MTAB-17049. All Python code is available at https://github.com/wade-lab/HppXcrod2.

## ACKNOWLEDGEMENTS

We thank the Wadsworth Center Applied Genomic Technologies Core Facility, and the Wadsworth Center Media and Glassware Core Facility for assistance. We thank José Luis Puente for generously providing the JPCR7 and JPCR9 strains of *C. rodentium*. We thank Siân Owen, Thomas Bartlett, Irina Artsimovitch, and David Grainger for helpful discussions.

---

**Figure S1.**
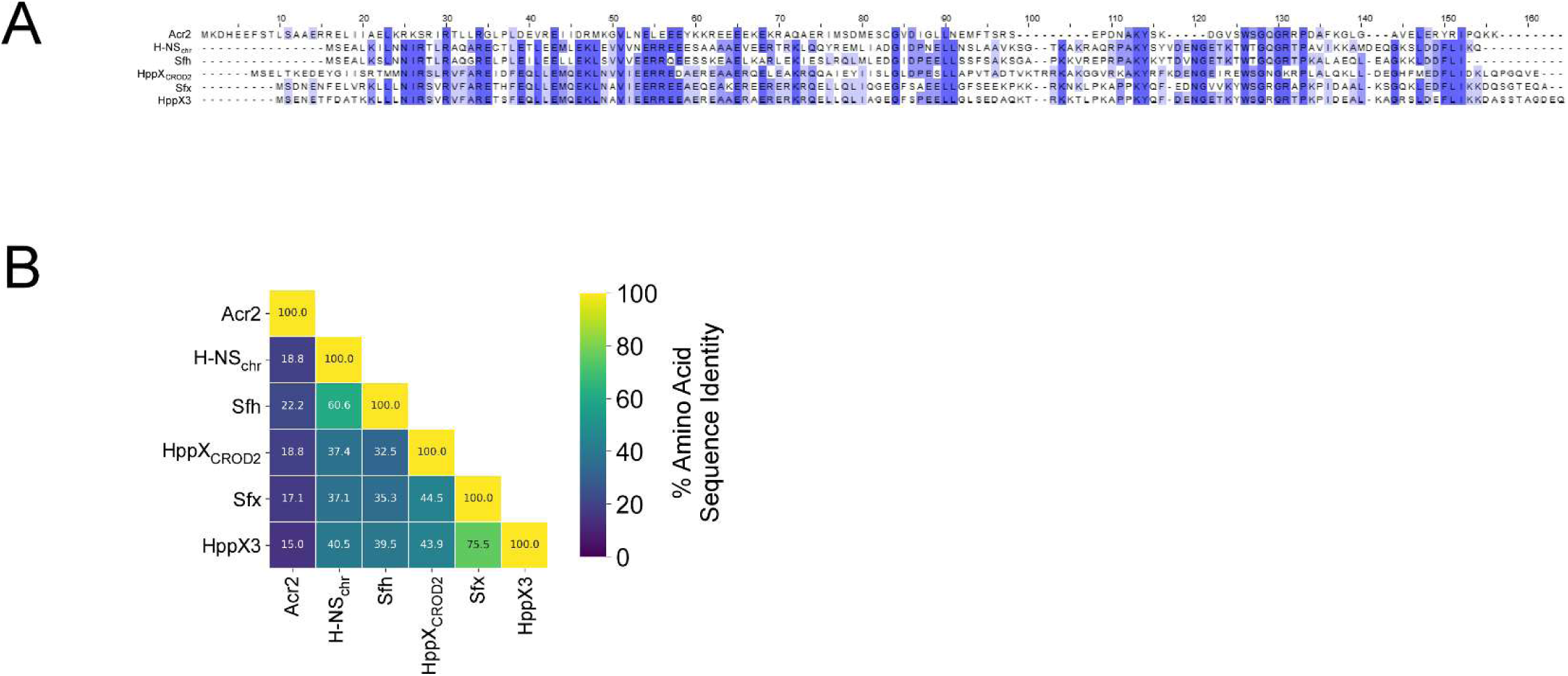
Sequence comparison of H-NS_chr_ and Hpp proteins. **(A)** Sequence alignment of H-NS_chr_ (from *C. rodentium*) and five Hpp proteins: HppX_CROD2_, Sfx, Sfh, HppX3, and Acr2. Conserved residues are shaded, with darker shading indicating broader conservation. **(B)** Heatmap showing the percent amino acid sequence identity for pairwise comparisons between HppX_CROD2_, Sfx, Sfh, HppX3, and Acr2.

**Figure S2.**
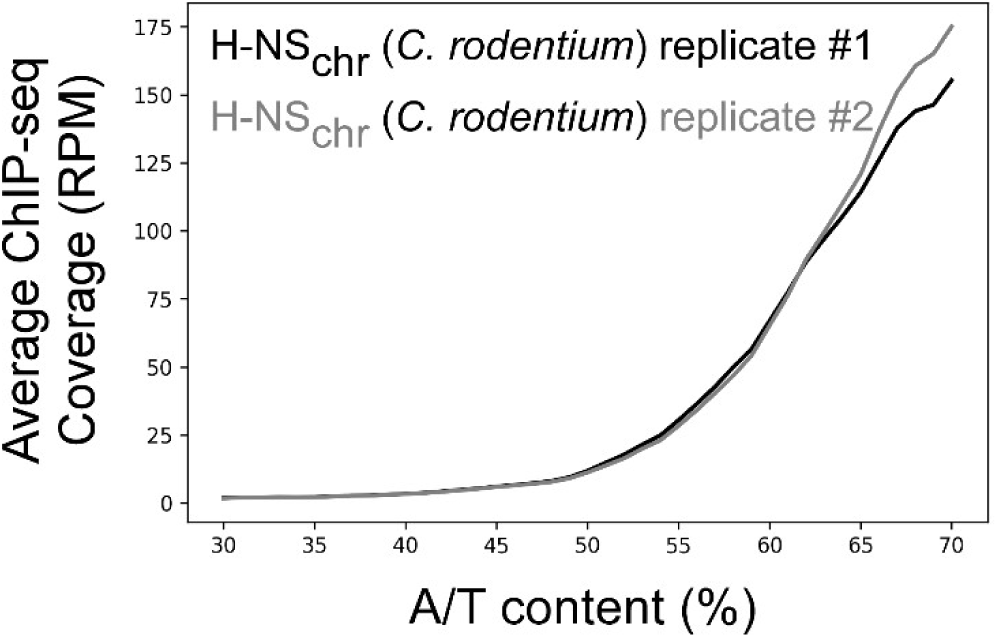
H-NS_chr_ associates with A/T-rich regions of the *C. rodentium* chromosome. A/T content (%) and ChIP-seq occupancy (reads per million) for H-NS_chr_ were determined for 1,000,000 randomly selected 101 bp regions. Mean ChIP-seq occupancy is shown for bins of A/T content that span 1% each.

**Figure S3.**
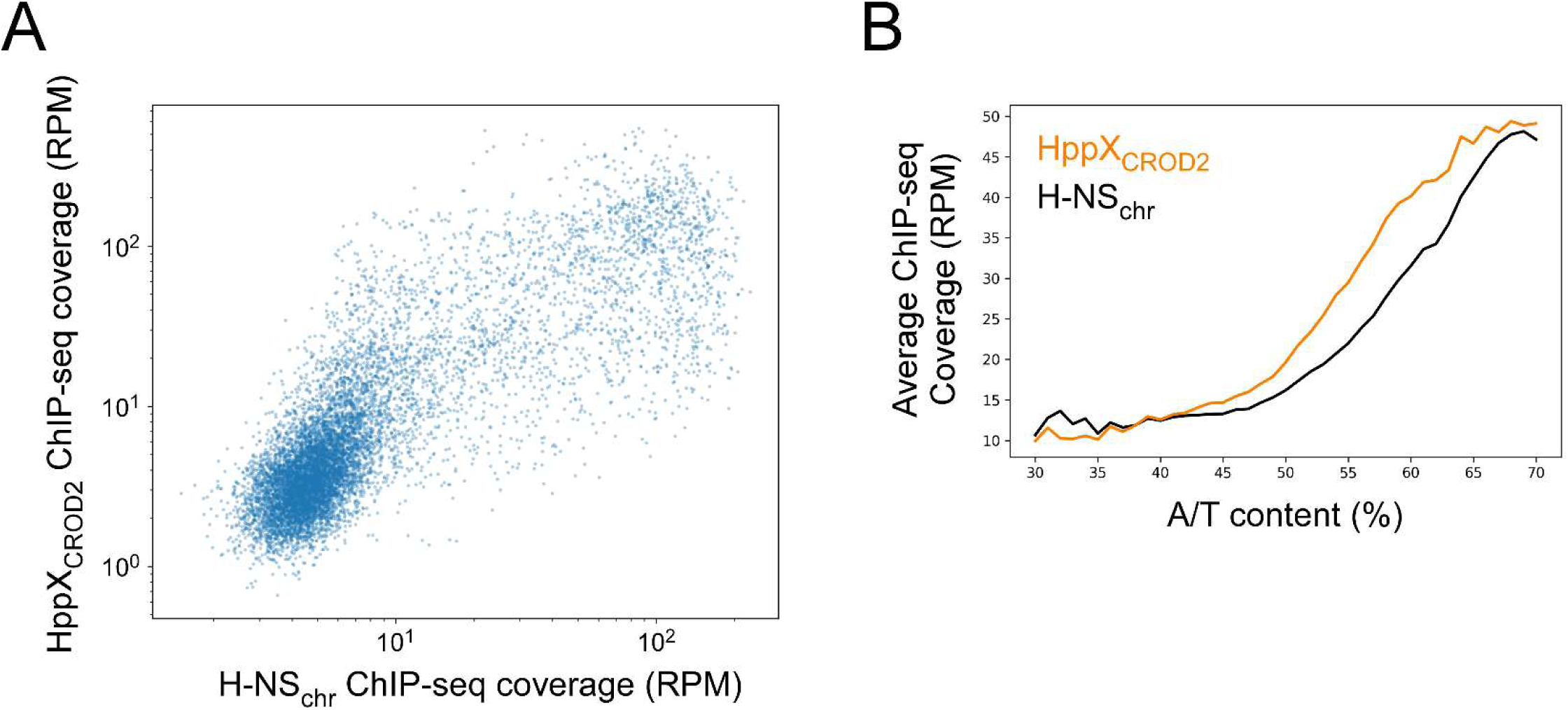
Comparison of H-NS_chr_ and HppX_CROD2_ binding to the *E. coli* chromosome. **(A)** Scatter-plot showing H-NS_chr_ (*x*-axis) and HppX_CROD2_ (*y*-axis) ChIP-seq occupancy (reads per million) at 10,000 randomly selected 100 bp regions of the *E. coli* chromosome. One of two replicate datasets is shown for each experiment. **(B)** A/T content (%) and ChIP-seq occupancy (reads per million) for H-NS_chr_ (black line) and HppX_CROD2_ (orange line) were determined for 1,000,000 randomly selected 101 bp regions. Mean ChIP-seq occupancy is shown for bins of A/T content that span 1% each. One of two replicate datasets is shown for each experiment.

**Figure S4.**
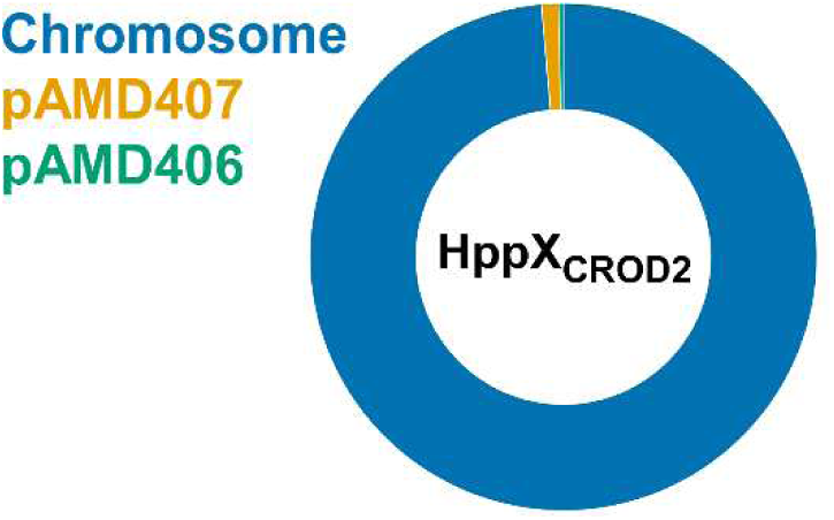
HppX_CROD2_ does not selectively bind any plasmid where it is encoded. Donut-chart showing the proportion of ChIP-seq coverage mapping to the *E. coli* chromosome and the pAMD407 and pAMD406 plasmids for HppX_CROD2_-FLAG_3_. Note that HppX_CROD2_-FLAG_3_ is encoded on pAMD406.

**Figure S5.**
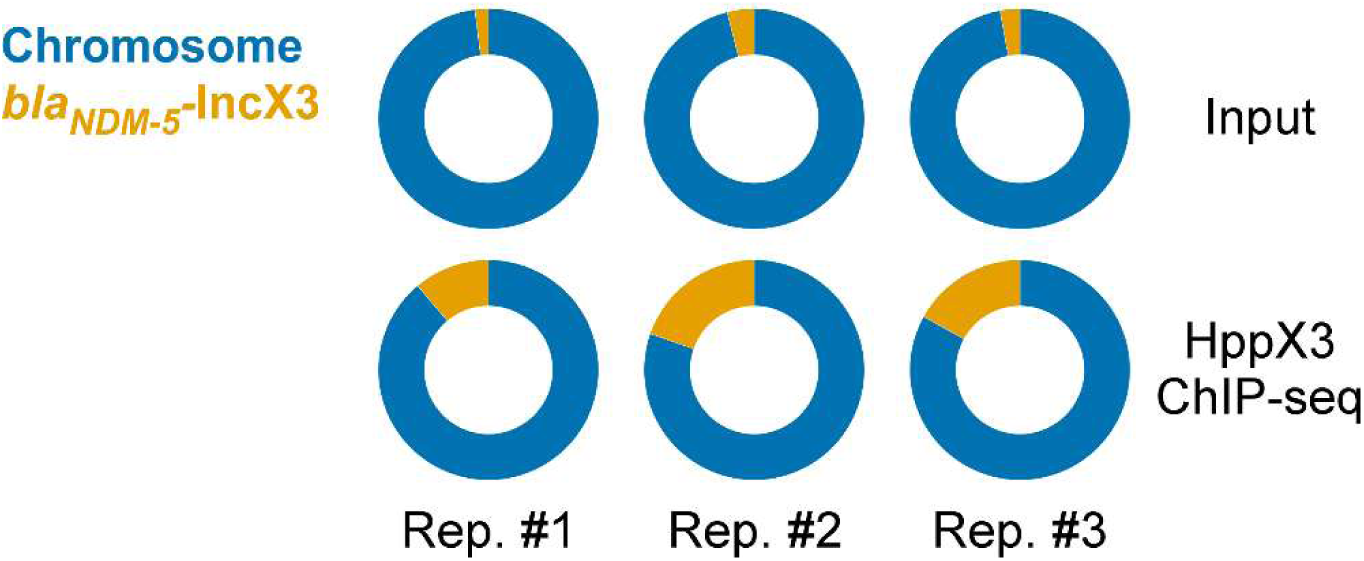
Selective binding of HppX3 to the *bla*_NDM-5_-IncX3 plasmid. Donut-charts showing the proportion of sequence read coverage mapping to the *E. coli* chromosome and the *bla*_NDM-5_-IncX3 plasmid for HppX3-FLAG ChIP-seq and corresponding input (i.e., pre-immunoprecipitation) samples (from (12)).

**Figure S6.**
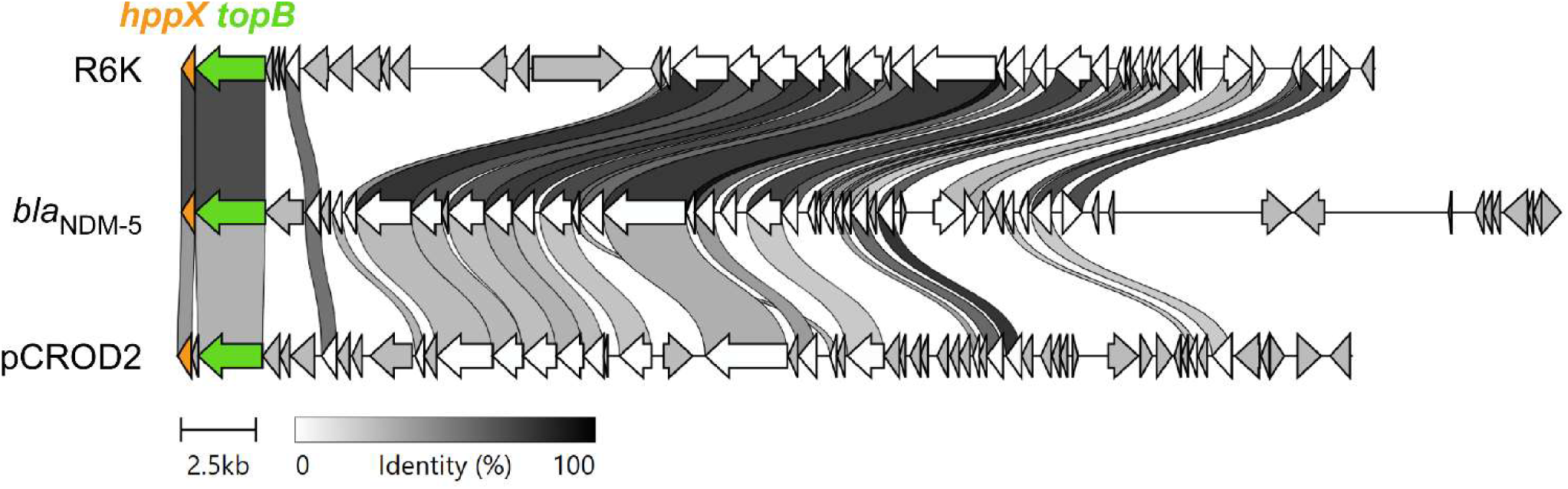
Synteny of *hns* and *topB* genes is shared across the R6K, *bla*_NDM-5_-IncX3, and pCROD2 plasmids. Gene-level alignment of the R6K, *bla*_NDM-5_-IncX3, and pCROD2 plasmids, indicating homologous genes (linked by gray shading; the darkness of the gray indicates the percent amino acid identity between encoded protein pairs). *hppX* and *topB* (encodes topoisomerase) genes are shown in orange and green, respectively.

